# Muscle-specific increased expression of *JAG1* improves skeletal muscle phenotype in dystrophin-deficient mice

**DOI:** 10.1101/2025.03.12.642857

**Authors:** Felipe de Souza Leite, Matthias R. Lambert, Tracy Yuanfan Zhang, James R. Conner, Joao A. Paulo, Sheldon Furtado Oliveira, Sanjukta Thakurta, Jennifer Bowles, Emanuela Gussoni, Steven P. Gygi, Jeffrey J. Widrick, Louis M. Kunkel

## Abstract

Therapeutic strategies for Duchenne Muscular Dystrophy (DMD) will likely require complementary approaches. One possibility is to explore genetic modifiers that improve muscle regeneration and function. The beneficial effects of the overexpression of Jagged-1 were described in escaper golden retriever muscular dystrophy (GRMD) dogs that had a near-normal life and validated in dystrophin-deficient zebrafish (1). To clarify the underlying biology of *JAG1* overexpression in dystrophic muscles, we generated a transgenic mouse (mdx^5cv^-*JAG1*) model that lacks dystrophin and overexpresses human *JAG1* in striated muscles. Skeletal muscles from mdx^5cv^-*JAG1* and mdx^5cv^ mice were studied at one, four, and twelve-month time points. *JAG1* expression in mdx^5cv^-*JAG1* increased by three to five times compared to mdx^5cv^. Consequently, mdx^5cv^-*JAG1* muscles were significantly bigger and stronger than dystrophic controls, along with an increased number of myofibers. Proteomics data show increased dysferlin in mdx^5cv^-*JAG1* muscles and an association of Nsd1 with the phenotype. Our data supports the positive effect of *JAG1* overexpression in dystrophic muscles.

**Significance Statement:** Duchenne Muscular Dystrophy (DMD) patients present a progressive decline in motor function. DMD is caused by mutations in the *DMD* gene that lead to the absence of dystrophin - an essential component of muscle cells. However, dystrophin-deficient dogs overexpressing JAG1 had a normal lifespan with remarkable motor function. In this study, we increased expression of human *JAG1* in mouse skeletal muscles lacking dystrophin to explore mechanisms responsible for these benefits. Our observations show that overexpression of *JAG1* counterbalances the lack of dystrophin by generating bigger and stronger muscles as the mouse ages. Moreover, our proteomics dataset suggests a role of dysferlin in the phenotype. Therefore, our study supports the exploration of *JAG1* in pre-clinical models.

## Introduction

Duchenne Muscular Dystrophy (DMD) is an X-linked recessive neuromuscular disease with devastating consequences. It is caused by frameshift or truncating mutations in the *DMD* gene (2), leading to the absence of dystrophin in striated muscles (3). DMD incidence is 1 in 3,500 to 5,000 male births (4). Initial symptoms arise between three to five years of age when patients present with constant falls and proximal muscle weakness. As the disease progresses, distal muscles are also weakened; ten- to twelve-year-old patients become wheelchair dependent and may need ventilatory assistance. The leading causes of death are respiratory insufficiency and cardiomyopathy, and the current median lifespan for patients undergoing corticosteroid treatment is approximately 28 years (5).

Heterogeneity observed among patients’ disease progression and among animal models led to the discovery of a handful of genetic modifiers. The rare G allele in the promoter of *SPP1* (rs28357094) correlates with rapid progression and greater severity (6). LTBP4 was initially found in a DMD mouse model (7), and LTBP4 inhibition with a monoclonal antibody showed beneficial effects in mouse muscles (8). In human patients, the *LTBP4* IAMM haplotype in homozygosis was associated with prolonged ambulation (9). The common null polymorphism R577X in *ACTN3* leads to reduced power outcomes and a longer time to complete a 10-meter walking test in DMD patients. Consistently, α-actinin-3 deficiency in the dystrophin-deficient mouse model (mdx) led to reduced muscle strength compared to mdx alone, yet aged double knockout mice displayed phenotypic improvements, as noticed by improved histological features and more significant muscle recovery from fatigue (10). Recently, a genome-wide association study using 419 DMD patients found six additional modifier candidates (11). Finally, myostatin inhibition in a DMD murine model showed partial improvements (12). These observations show the complex mechanisms underlying DMD progression and raise the possibility for modifier-inspired treatments (13).

The Notch pathway is a conserved cell-to-cell communication system that regulates cell-fate specification, differentiation, and patterning. Several studies support the role of Notch modulation in muscle health. Jagged-1 is a canonical ligand in the Notch pathway, and its overexpression was found in two golden retriever muscle dystrophy (GRMD) dogs that largely escaped their expected disease progression (1, 14, 15). A Notch3^-/-^ mouse model exhibited an overgrown muscle phenotype after repetitive injuries (16). Constitutively active Notch1 expression under muscle creatine kinase (MCK) in mouse myofibers showed improved stem cell niche and regeneration (17). Hyperactivation of the Notch pathway in a mouse model of neurofibromatosis type 1 is associated with impaired function of muscle progenitors, and early postnatal inhibition of the Notch pathway showed beneficial effects (18). In humans, a reduction in Notch signaling due to a missense mutation in POGLUT1 leads to a type of autosomal recessive limb-girdle muscular dystrophy (19). Finally, a whole-exome investigation of an international cohort of 23 patients with unexplained muscular dystrophy linked their disorder to rare *JAG2* variants that caused a novel form of autosomal recessive muscular dystrophy (20). Thus, Notch signaling modulation may generate beneficial or harmful effects in skeletal muscle.

The finding that jagged-1 was overexpressed in escaper GRMD dogs established the grounds for a therapeutic exploration of the Notch pathway in dystrophin-deficient muscles, bypassing the rescue of dystrophin expression (1). However, a beneficial effect of a putative therapy relies on clarifying its underlying biology. Thus, we generated a mouse model to study the consequences of *JAG1* overexpression in dystrophic skeletal muscles. The mdx^5cv^ mouse has a mutation in exon 10 that disrupts full-length dystrophin expression; it is more affected than the standard mdx and has fewer revertant fibers (21). This study used a mdx^5cv^-*JAG1* mouse model, a dystrophin-null mdx^5cv^ mutant that specifically overexpresses human *JAG1* in striated muscles. Our data supports that a balanced overexpression of human *JAG1* in dystrophic muscles can counterbalance disease progression by increasing muscle mass and force over time. Additionally, by proteomics screening we identified increased expression of dysferlin and Nsd1 in mdx^5cv^-*JAG1* muscles, suggesting an association among jagged-1, dysferlin, and Nsd1.

## Results

### mdx^5cv^-*JAG1*’s muscles overexpress *JAG1* throughout life

We took advantage of three previously described mouse models to increase *JAG1* expression in the striated muscles of the mdx^5cv^ mouse. The R26-LSL-*JAG1* (22) carrying the human *JAG1* cDNA located immediately 3’ of a *loxP*-flanked STOP cassette that blocks transcription, and the MCK-Cre (23) mouse that allows Cre-mediated recombination in striated muscles **(Figure 1A)**. Thus, the mdx^5cv^ was crossed with the R26-LSL-*JAG1* line to insert the human *JAG1* cDNA. After at least two generations, female mdx^5cv^;R26-LSL-*JAG1*^tg/tg^ were crossed with the MCK-Cre line, allowing tissue-specific expression of human *JAG1* **(Figure 1B)**. We studied *JAG1* heterozygotes since the escaper dogs were heterozygotes for the variant in the *JAG1* promoter (1) and to avoid possible toxic effects. Our experimental animal groups were males, either mdx^5cv^;R26-LSL-JAG1^Tg/WT^;MCK-Cre or mdx^5cv^;R26-LSL-JAG1^Tg/W^, respectively named mdx^5cv^-*JAG1* and mdx^5cv^.

**Figure 1.**
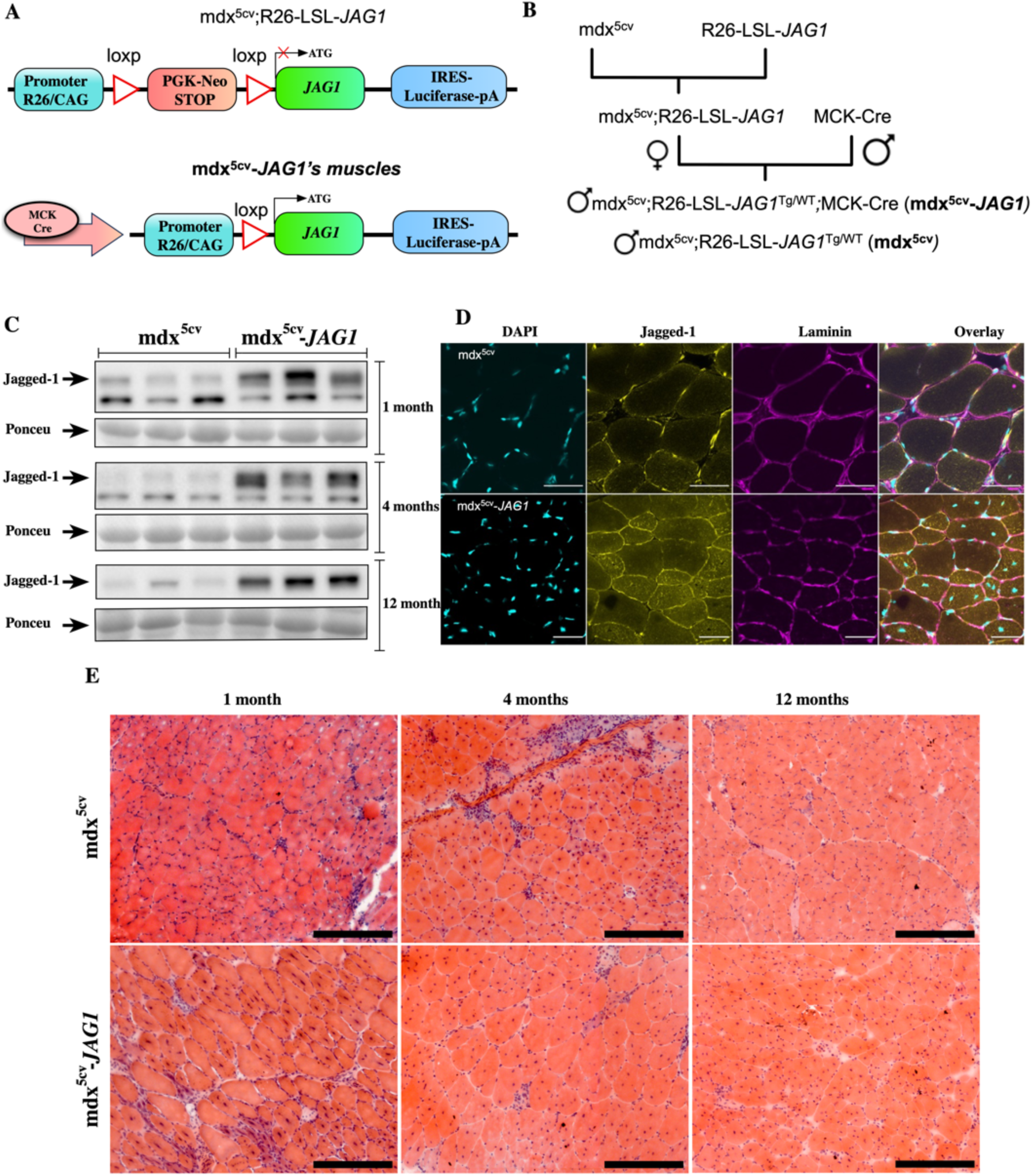
mdx^5cv^-*JAG1* mice overexpress Jagged-1 across life. **(A)** The top portion shows the construct used for our mdx^5cv^ controls, which had human *JAG1* transcription blocked by a *loxP*-flanked STOP cassette. The bottom portion shows the final construct in which muscle creatine kinase drives the expression of Cre recombinase and allows overexpression of *JAG1* and luciferase. **(B)** Schematic diagram of the breeding strategy to generate mice with striated muscle-specific Jagged-1 overexpression. Mdx^5cv^;R26-LSL-*JAG1* mice crossed with MCK-Cre mice will produce Mdx^5cv^;R26-LSL-*JAG1*^Tg/WT^,MCK-Cre offspring, which overexpress Jagged-1 in skeletal and cardiac muscle specific. **(C)** Western blot images showed increased expression of Jagged-1 at the three time points tested. Intensity changes were calculated using the Ponceau as loading control for each timepoint. All original membranes are in Supplemental Figure 1. **(D)** Immunofluorescence images of TA cross-sections at four months stained for DAPI (Cyan), jagged-1 (Yellow), and laminin (Magenta). **(E)** Hematoxylin and Eosin (H&E) staining at the three timepoints tested (Columns). The top and bottom rows show images from mdx^5cv^ and mdx^5cv^-*JAG1*, respectively. Scale bars are 250 μm.

To comprehensively understand the effects of *JAG1* overexpression in dystrophic muscles, we performed non-survival experiments in mice at one, four, and twelve months. For these three time points, western blots using protein extracts from quadriceps muscles confirmed increased jagged-1 expression **(Figure 1C and Supplemental Figure 1)**. Consistently, immunostaining of tibialis anterior (TA) muscle sections at four months reinforced our results as increased jagged-1 staining in the mdx^5cv^-*JAG1* muscles was observed, particularly at the sarcolemma **(Figure 1D)**. Thus, our data confirmed increased jagged-1 protein in skeletal muscles during the timeframe pre-defined for this study.

Hematoxylin and eosin (H&E) staining of TA whole sections showed central nuclei at all time points in both groups **(Figure 1E)**. By the first month, muscles already displayed dystrophic features, but mdx^5cv^-*JAG1* had a higher density of possible regenerative fiber pockets. At four months, mdx^5cv^ had extensive inflammation areas and scarring **(Figure 1E)**; while these features were also present in the mdx^5cv^-*JAG1*, they were less extensive. Sections from one-year-old animals in both groups were similar, albeit mdx^5cv^-*JAG1* muscle sections had a larger cross-sectional area than controls.

### The mdx^5cv^-*JAG1*’s skeletal muscles are bigger than the mdx^5cv^

One-month-old mdx^5cv^-*JAG1* and mdx^5cv^ had similar body and muscle mass **(Figure 2A)**. While body mass was still remarkably similar between both groups at the four-month-old time point, the average mass of mdx^5cv^-*JAG1* quadriceps muscles were ≈ 15% greater, thereby comprising a significantly higher portion of their body mass **(Figure 2B and C)**. Four-month-old mdx^5cv^-*JAG1*’s TA muscles were not significantly heavier (**Figures 2D and E**).

**Figure 2.**
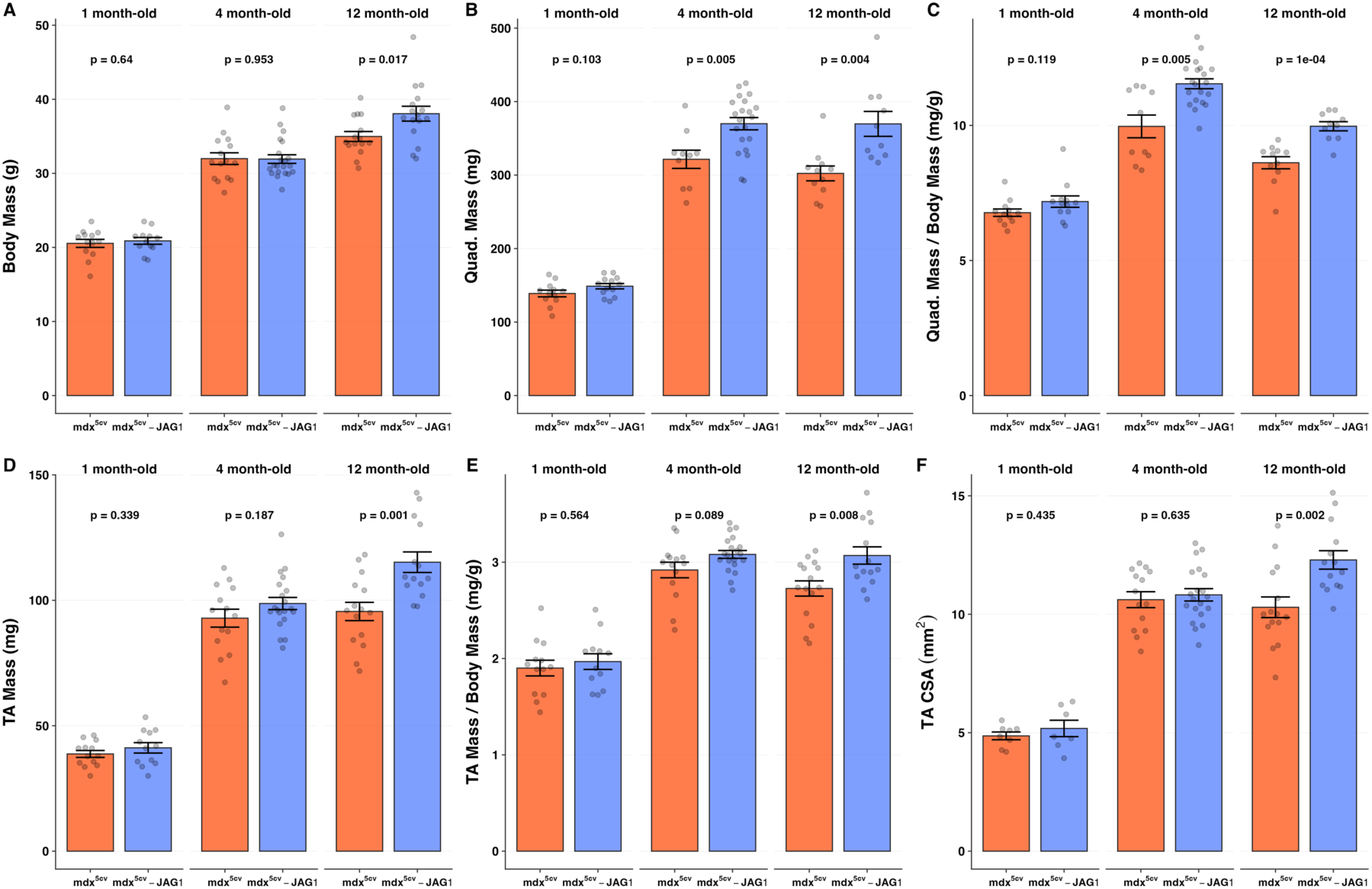
Changes in body and muscle mass over time. **(A -F)** Plots for whole body mass, dissected quadriceps mass, quadriceps mass normalized per body mass, dissected tibialis anterior (TA) mass, TA mass normalized per body mass, and TA cross-section area for one-, four, and twelve months. The mdx^5cv^ and mdx^5cv^-*JAG1* groups are in red and blue, respectively. Plots show mean ±SEM and p-value. Each gray dot represents data collected from one animal.

Differences increased further by one year when the mdx^5cv^-*JAG1* animals were ≈ 9% heavier **(Figure 2B)**, and their TA and quadriceps mass were ≈ 20% greater than mdx^5cv^ **(Figure 2B and D)**. Again, skeletal muscles comprised more of mdx^5cv^-*JAG1* total body mass than mdx^5cv^ **(Figure 2C and E)**. Consistently, mdx^5cv^-*JAG1* TA cross-section area (CSA) was 19.4% larger than mdx^5cv^ at 12 months and increased by ≈ 13% compared to four-month-old mdx^5cv^-*JAG1*, emphasizing that mdx^5cv^-*JAG1* muscle mass continued to grow. In contrast, the average CSA of mdx^5cv^’s TA muscles peaked at four months **(Figure 2F)**. To sum up, our data shows that *JAG1* overexpression has a mild effect on dystrophic muscle histology but suggests an increase in regenerative areas at one month, followed by greater muscle mass later in life.

### Muscles from mdx^5cv^-*JAG1* change their myofiber profile with time

We investigated changes in the size and fiber type composition of mdx^5cv^-*JAG1* muscles using whole sections of TA muscles obtained from the thickest part of each muscle **(Figure 3A and Supplemental Figure 2)**. In the first month, analyses of the myofiber minimum Feret diameter with laminin staining showed a similar density distribution across the examined range between groups (10 μm to 100 μm). However, the mdx^5cv^-*JAG1* median was slightly lower due to the higher density of fibers smaller than ≈ 20 μm **(Figure 3B)**. At the first time point, both groups show the narrowest distribution of fiber dimensions **(Figure 3C)**.

**Figure 3.**
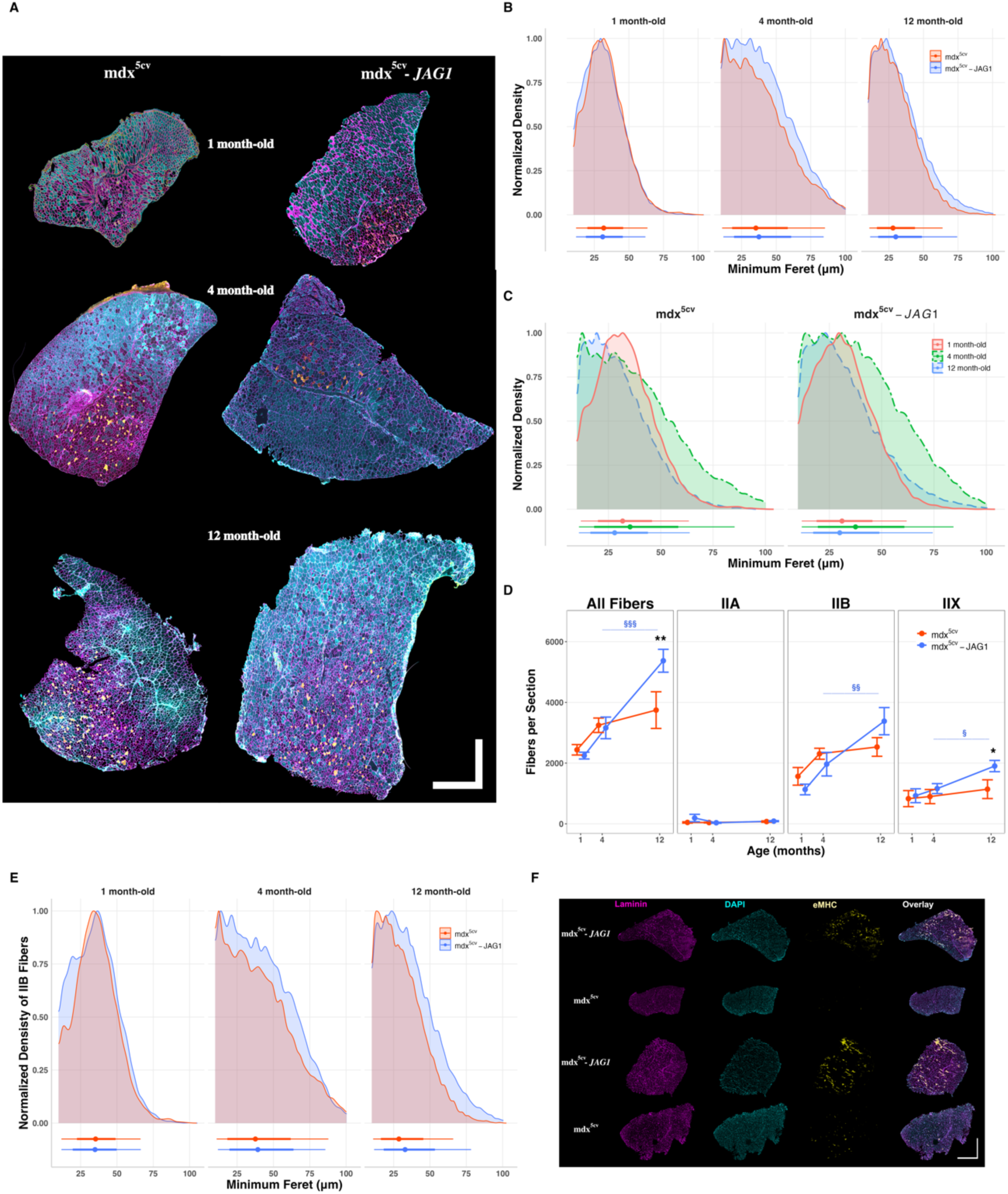
mdx^5cv^-*JAG1*’s sections were bigger and showed differences in fiber type and size. **(A)** Overlay of stained sections obtained from the thickest portion of the TA muscle. The left and right rows show sections for mdx^5cv^ and mdx^5cv^-*JAG1* at one-, four-, and twelve-month timepoints. Sections were stained for laminin (Magenta), myosin heavy chain type I, type IIA (yellow), and IIB (Cyan). Scale bars are 1000 μm**. (B)** Histograms comparing the normalized density of fibers from 10 μm to 100 μm between mdx^5cv^ (red) and mdx^5cv^-*JAG1* (blue). **(C)** Same as the figure 3B but organized by genotype to show the temporal progression of fiber density per minimum Feret. **(D)** Comparison of the total number of fibers per section at the three timepoints, and the number of fibers of each fiber type for all muscle sections from mdx^5cv^ and mdx^5cv^-*JAG1* mice. Type I fibers were excluded since they are less than 1% of the fiber type composition. Not stained fibers were considered type IIX. **(E)** Histograms comparing the normalized density of type IIB fibers from 10 μm to 100 μm. **(F)** mdx^5cv^-*JAG1* mice have an increased number of newly formed fibers at one month. From left to right, the figure shows laminin staining in magenta, nuclear DNA staining with DAPI in cyan, embryonic myosin heavy chain (eMyHC) in yellow, and the overlay of all channels. From top to bottom, the figure intercalates two different mice from each group, i.e. mdx^5cv^-*JAG1* and mdx^5cv^. The scale bar is 500 μm. Additional eMyHC from the other four- and twelve-month-old time points are shown in Supplemental Figure 3. Error bars are SEM and level of significance between genotypes in the same age is indicated as * = p < 0.05 and ** = p < 0.01. Level of significance between time points of the same genotype is indicated as ^§^ = p < 0.05, ^§§^ = p < 0.01, and ^§§§^ = p < 0.001 – the blue color indicates the mdx^5cv^-*JAG1* genotype, no difference was observed in the mdx^5cv^ genotype.

Groups profiles diverge at four months. Overall fiber hypertrophy from one to four months and the higher density of small fibers drives a broader distribution in both groups – the widest observed **(Figure 3C)**. However, the distribution of the mdx^5cv^-*JAG1* group is broader, with the most frequent quartile of fiber distribution ranging from 10 μm to 50 μm, whereas controls range from 10 μm to ≈ 40 μm. Thus, the median minimum Feret of both groups shifts to higher diameter relative to the first-month distribution, with a more significant displacement for the mdx^5cv^-*JAG1*, surpassing control **(Figures 3B and C)**.

As the mice age to one year old, both groups show a narrower distribution profile than at four months **(Figure 3C)**. The profiles of both groups have similar shapes, but the mdx^5cv^-*JAG1* distribution is shifted to the right, still showing a higher density of larger fibers and a higher median value **(Figures 3B and C)**. Additionally, the absolute number of fibers per section analyzed was 51% greater in the mdx^5cv^-*JAG1* at the one-year time point **(Figure 3D)**, a significant increase from four months to one-year-old time point, which was not observed in the mdx^5cv^ group **(Figure 3D)**.

Concomitantly, sections were stained for myosin heavy chain type I, IIA, IIB, and laminin. **(Figure 3A)**. At one-month-old, the sections of mdx^5cv^-*JAG1* showed ≈ 32% fewer type IIB fibers than mdx^5cv^, while IIX percentages were similar. The average number of fibers type IIB and IIX per cross-section increases in both groups from the first month to the fourth. However, mdx^5cv^-*JAG1* displays a steep increase in the twelve-month time point, while the mdx^5cv^ group remains stable **(Figure 3D)**. The density distribution of IIB fibers shows a particular deviation from controls at all time points. In the first timepoint, mdx^5cv^-*JAG1* mice have a greater density of small IIB fibers (10 μm to 25 μm), but on the other timepoints, mdx^5cv^-*JAG1* display greater densities along most of the range **(Figure 3E)**.

To further address the potential increase in newly formed fibers, sections were stained with anti-embryonic myosin heavy chain (eMyHC). One-month mdx^5cv^-*JAG1* sections presented more and larger eMyHC fiber clusters than mdx^5cv^ **(Figure 3F)**. Increased staining was still observed in the four and twelve-month-old time points, albeit fewer eMyHC-expressing fibers could be observed at these time points **(Supplemental Figure 3)**. In summary, our data indicates that an increased number of newly formed fibers in the mdx^5cv^-*JAG1* group and a sharp increase in type IIB and IIX fibers drive the continuous increase in muscle size observed in mdx^5cv^-*JAG1* mice.

### Overexpression of *JAG1* enhances skeletal muscle force

We investigated changes in force using TA muscles *in situ*. Four-month-old mdx^5cv^-*JAG1* muscles generated stronger peak forces than mdx^5cv^ **(Figure 4A),** and normalization per CSA showed higher specific force **(Figure 4B)**. Next, muscles underwent a force-frequency protocol with trains of electrical impulses ranging from 20 to 200 Hz with one minute resting interval between each train **(Figure 4C)**. Consistent with our muscle mass findings, the first time point at one month did not show an increase in force, but four- and twelve-month-old mdx^5cv^-*JAG1* generated significantly higher forces than mdx^5cv^ at several frequencies (**Figure 4D)**.

**Figure 4.**
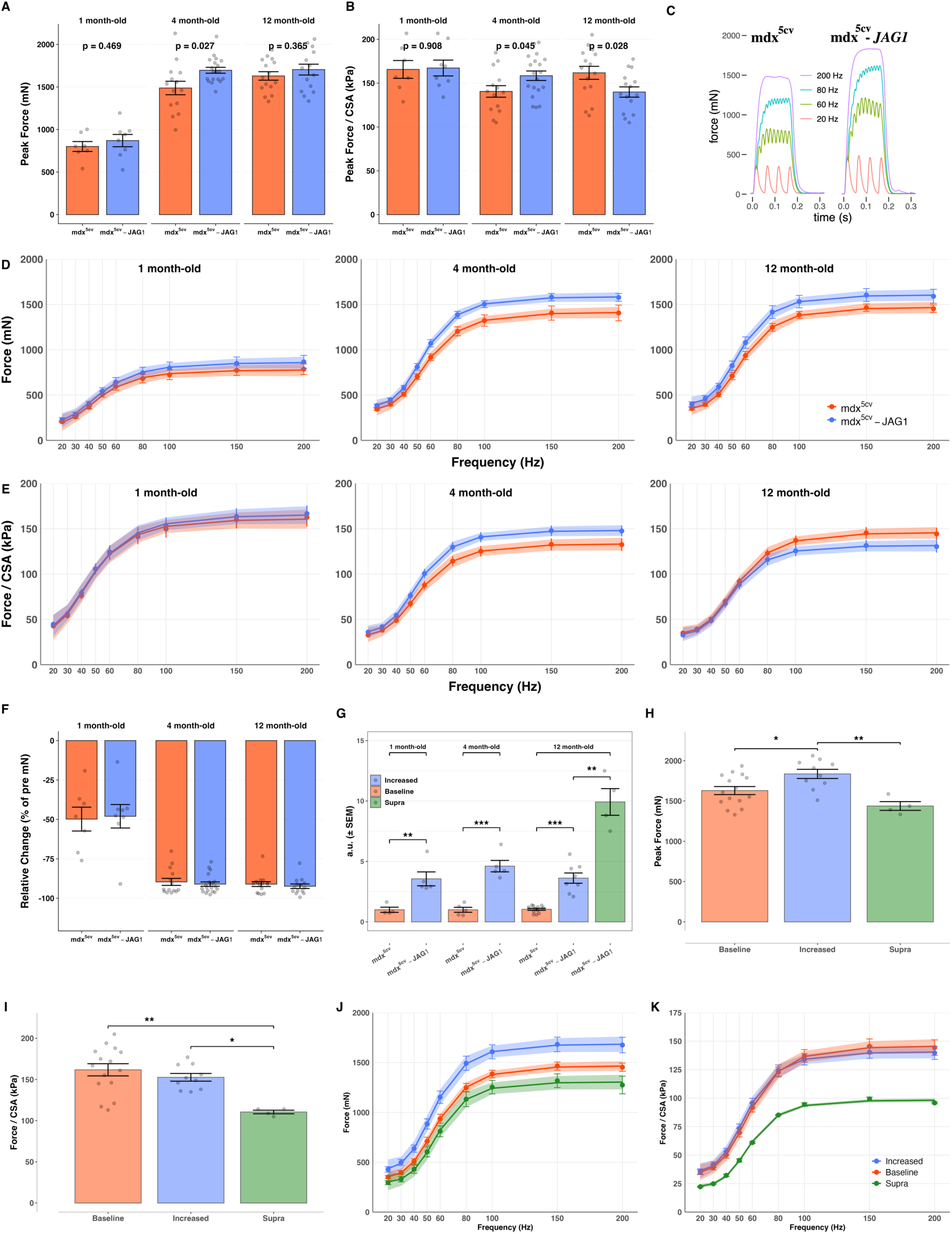
Overexpression of Jagged-1 increases whole-muscle force over time. **(A)** Peak force of TA muscles from each animal per time point. **(B)** Specific force (Peak force normalized by cross-section area (CSA)). **(C)** Representative force records of both genotypes tested. **(D)** Absolute force of TA muscles vs. frequency of stimulation. Error bars represent SEM and the shaded area indicates the estimated 95th confidence interval. For each curve, an F-test was used to compare a genotype-specific fitting model against a shared model across genotypes. A genotype-specific model provided the best fit for data at the four- and twelve-month time points (F(4,289) = 12.47, p < 0.001 and F(4,253) = 7.71, p < 0.0001, respectively), but not at the one-month time point (F(4,124) = 1.66, p = 0.163). **(E)** Similar to D, but for specific force. A genotype-specific model provided the best fit for data at the four- and twelve-month time points (F(4,289) = 7.00, p < 0.0001 and F(4,253) = 3.43, p = 0.0095, respectively), but not at the one-month time point (F(4,124) = 0.16, p = 0.959). **(F)** Change in isometric force of TA muscles after the eccentric contraction protocol. Values are the post-protocol isometric force expressed relative to the pre-protocol isometric force. Supplemental Figure 4 shows the progressive decline in force obtained during these experiments. **(G)** Jagged-1 expression in quadriceps muscles assessed by Western blot. Intensity of chemiluminescence was normalized to whole protein lysate content (i.e. Ponceau staining intensity). Each dot represents Western blot analysis performed in an individual mouse. Large discrepancies were observed within the mdx^5cv^-Jag1 group at 12 months of age in which most mice display increased Jagged-1 expression as similarly seen at 1 and 4 months of age (3- 5-fold increase), while a small cohort of mice displays 10-fold increased expression (referred to as supra group). **(H-K)** Peak force, specific force, force-frequency, and specific force-frequency relationships when animals with supra-abundance of Jagged-1 are considered as separate group. A two-way ANOVA showed a significant effect for level of jagged-1 abundance in Force and Force/CSA (p = 0.000958 and p = 0.00435 respectively). Bonferroni-adjusted pairwise comparing Force (Fig. H) showed significant differences between Baseline vs. Increased (p.adj= 0.0284) and Increased vs. Supra (p.adj= 0.00288), whereas Baseline vs. Supra was not significant (p.adj= 0.213). The same statistical test for Force/CSA (Fig. I) showed differences between Baseline vs. Supra (p.adj= 0.00142) and Increased vs. Supra (p.adj= 0.0131), whereas Baseline vs. Increased was not significant (p.adj= 1.000). Error bars are SEM and shaded area is the estimated 95^th^ confidence interval, and level of significance is indicated as * = p < 0.05, ** = p < 0.01, and *** = p < 0.001.

Dystrophin-deficient muscles are sensitive to damage when actively stretched (eccentric contractions). Muscles from mdx^5cv^-*JAG1* did not show an improvement in resistance to stretch, i.e., while their muscles generated higher forces before the stretch-induced injury protocol, their relative force decay was not different from mdx^5cv^ controls **(Figure 4F and Supplemental Figure 4)**.

At first glance, the experiments described above showed an unexpected specific force profile at twelve months. While one-year-old mdx^5cv^-*JAG1*’s muscles were stronger, their specific force was lower than controls **(Figure 4B and E)**. However, during our study, we noticed a specific set of siblings mdx^5cv^-*JAG1* mice that had a much higher abundance of jagged-1 **(Figure 4F)**. These four animals generated significantly less force and specific force than the other 11 mdx^5cv^-*JAG1* mice studied and were primarily responsible for the unexpected drastic reduction in force and, particularly, specific force **(Figures 4 G-I)**. In conclusion, from an *in situ* physiology perspective, mdx^5cv^-*JAG1* mice muscles generate more force but are not more resistant to actively stretch-induced damage. Nevertheless, it is important to consider our circumstantial observation of mice expressing supra levels of jagged-1 - suggesting toxicity. Of note, these four animals were only used in physiology experiments.

### Increased Dysferlin and Nsd1 in mdx^5cv^-*JAG1* muscles

We conducted global proteomics using quadriceps muscles collected at all time points to begin exploring the mechanisms behind the mdx^5cv^-*JAG1* phenotype. Dysferlin, a membrane-associated protein responsible for sarcolemma repair (24), had ≈ 60% increased abundance at the first and third time point **(Figures 5A-C)**. Notably, two out of three four-month-old samples had higher levels of dysferlin than mdx^5cv^ animals. Thereby, eight out of nine mdx^5cv^-*JAG1* samples had higher dysferlin expression than their respective time point mdx^5cv^ controls **(Figure 5D)**.

**Figure 5.**
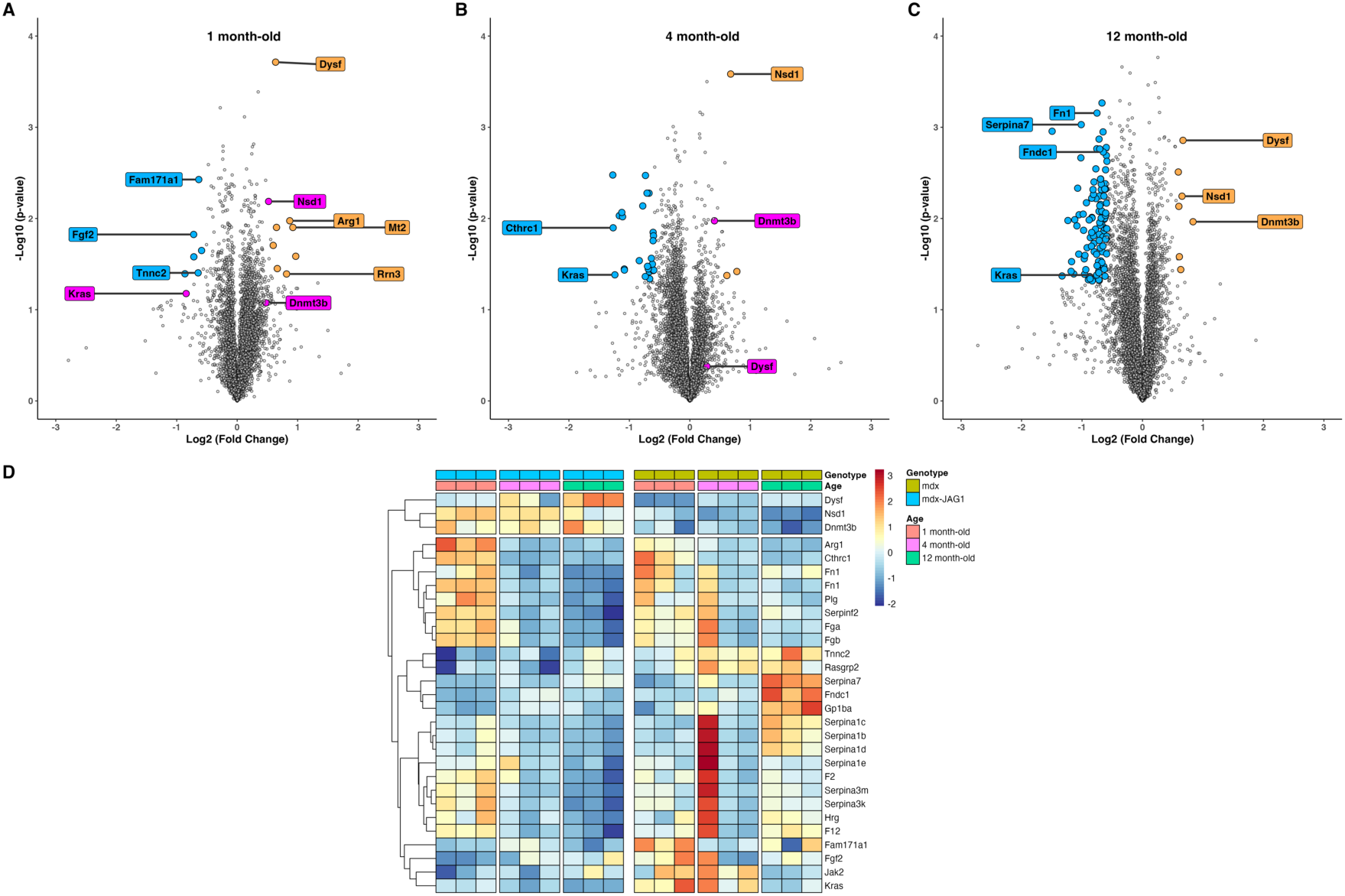
Proteomics using quadriceps muscles dissected from all the time points studied. **(A-C)** Each point in the volcano plot show the fold change of a protein comparing mdx^5cv^-*JAG1* over mdx^5cv^. Orange and blue dots show all proteins with statistically significant changes in expression and with fold change > 1.5. Magenta points show specific cases described in the main text. **(D)** Heat map showing the Z-score normalized per line of selected important in the context of this study.

Nds1, a histone methyltransferase that binds close to promoters of various genes with important roles in cell growth, cancer, and bone morphogenesis (25, 26) was detected among proteins with higher abundance and statistical significance. Dnmt3b was significantly increased in one-year-old mdx^5cv^-*JAG1* muscles **(Figures 5A-C)**. Dnmt3b is a *de novo* DNA methyltransferase that preferentially methylates gene bodies of actively transcribed genes (27). Haploinsufficiency of Nsd1 is associated with overgrown conditions, Sotos and Beckwith-Wiedemann syndromes (28), while mutations in the Dnmt3b are associated with forms of Facioscapulohumeral muscular dystrophy (28–30).

The number of differentially abundant (DA) proteins increased with age, supporting the temporal change in phenotype **(Figures 5A-C and Supplemental Figure 5)**. The first time point showed a set of 14 DA proteins **(Supplemental Table 1)**. Particularly, Arg1, Fam171a1, Fgf2, Mt2, Rrn3, and Tnnc2 have described roles in skeletal muscles. Gene set enrichment analysis (GSEA) suggested suppression of muscle-related and aerobic respiration pathways, while terms related to chromosome organization, chromatin remodeling, and immune system response were activated **(Supplemental Figures 6)**. The second time point analysis showed 28 DA proteins - 25 with lower abundance **(Supplemental Table 2)**. Among them are Cthrc1, Jak2, and Mylpf. Most of the top activated pathways relate to metabolism, including aerobic respiration and oxidative phosphorylation **(Supplemental Figure 6)**.

Finally, the third time point revealed 111 DA proteins **(Supplemental Table 3)**. Among the 104 proteins with lower abundance in the mdx^5cv^-*JAG1* group were two fibronectins, Fn1 and Fndc1, and a series of proteins of the serpin family. Activated enriched pathways included skeletal muscle tissue development, actin filament organization, and myotube differentiation **(Supplemental Figure 6)**. Overexpression analysis using the DE proteins shows blood coagulation and fibrinolysis pathways **(Supplemental Figure 6)**. Of note, while DMD patients display a hypercoagulable state (31, 32), genes associated with these pathways are downregulated in the mdx^5cv^-*JAG1* group. STRING analyses predicted the association of jagged-1 with Nsd1 and Dnmt3b through Notch1. Jagged-1 and Dnmt3b show connections with Kras – a constantly downregulated protein **(Supplemental Figure 7),** and a link with a large cluster of downregulated proteins in the one-year-old dataset.

## Discussion

Our findings show that increased *JAG1* expression in skeletal muscles increases muscle mass and force over time. The mdx^5cv^-*JAG1* mice have some of the classical features of dystrophin-deficient muscles (15). For instance, histology of TA muscles showed the presence of central nuclei, inflammation areas, and scar tissue **(Figure 1E)**. However, one-month-old mdx^5cv^-*JAG1*’s muscles showed increased regenerative fibers **(Figures 1E and 5)**. Consequently, as the mdx^5cv^-*JAG1* mice age, they develop larger and stronger muscles, with a sharp increase in the number of type IIB and IIX fibers and type IIB fiber cross-sectional area from four months to one year **(Figure 3E)**. Therefore, the increase in muscle mass seems to counterbalance the progressive weakness of dystrophin-deficient muscles.

A recent study observed that transplanting myogenic progenitors overexpressing the Notch1 intracellular domain (NICD) into injured mouse muscles increased the number of type IIX fibers (33). Dystrophin-deficient fast fibers, particularly IIX in humans, show earlier degeneration than slow fibers (34). Additionally, dystrophin-deficient animal models show that muscles composed mainly of type II fibers are more impacted by the lack of dystrophin (35–37). Thus, our findings with the mdx^5cv^-*JAG1* are particularly relevant for DMD since the overexpression of *JAG1* may spare the most damage-susceptible dystrophin-deficient muscles and support a Notch-related fiber type specification.

The phenotypic response to the overexpression of *JAG1* may not follow a linear relationship **(Figures 4F-J)**. Our circumstantial observation of mdx^5cv^-*JAG1* mice with ≈ 10-fold increase in *JAG1* suggests that modulation of the Notch pathway must be well-adjusted. Supporting this, the two escaper GRMD dogs presented only approximately 2-fold increased expression of jagged1 (1), and various phenotypes have been described depending on the modulation of the Notch pathway (16, 17, 38). Finally, our data supports the role of jagged-1 in the escapers’ dogs, but the absence of improved resistance to stretch suggests that jagged-1 was not the only player. For instance, our group reported the downregulation of PITPNA in the escaper dogs (39) and demonstrated how PITPNA inhibition with PDE10A improves the muscular phenotype in dystrophic zebrafish (40).

Our findings resemble myostatin inhibition, showing increased muscle mass through hypertrophic and hyperplasic events (41). Additionally, a myostatin knockout mouse model also shows a higher proportion of type IIB fibers (42), and myostatin blockade with antibodies in the mdx mouse showed improved muscle function, stronger absolute force, and no difference in eccentric force resistance to eccentric contractions (12). However, our data supports higher specific force in four-month-old mdx^5cv^-*JAG1* muscles, which was not observed with myostatin-blocking studies **(Figures 4B and D)**. Thus, an appropriate level of *JAG1* overexpression can increase muscle mass while maintaining muscle quality.

Dysferlin has an essential role in muscle repair, and mutations in the *DYSF* gene cause autosomal recessive limb girdle muscular dystrophy type 2B (LGMD2B) and Miyoshi myopathy (43, 44). Moreover, the mdx phenotype worsens when dysferlin is lacking (45). Our proteomics dataset suggests increased dysferlin expression in mdx^5cv^-*JAG1* muscles **(Figure 5)**. Notably, an mdx mouse model expressing the constitutively active form of Notch1 under MCK (MCK-N1ICD-*mdx*) also showed between 50 - 70% increase in dysferlin expression (17). Interestingly, human *DYSF* overexpression in non-dystrophic mice also showed a dose-response effect, where 2-fold increase is well-tolerated while > 30-fold higher abundance leads to a dystrophic phenotype (46). Thus, it is likely that dysferlin expression is under the control of Notch signaling and that the improved muscular phenotypes observed in these models are partially related to a balanced higher abundance of dysferlin.

Our data suggests an association with Nsd1, Dnmt3b, and Kras **(Figures 5D and E)**. Nsd3, a Nsd1 paralogue, has been associated with Notch pathway modulation (47), reinforcing an actual role of Nsd1 in the mdx^5cv^-*JAG1* phenotype. To cite additional proteomics findings, first, the upregulation of Rrn3 is associated with muscle hypertrophy and adaptation to exercise training (48). Second, Fgf2 has important roles in skeletal muscle physiology and pathology (49–51). Finally, there is a lower abundance of Fn1 and Ctrhrc1; the former is a DMD biomarker (52), and the latter was found to be increased in mdx muscles (53).

This is the first study using the mdx^5cv^-*JAG1* model, and future studies will address current limitations. First, future studies must explore cardiac function outcomes. Second, it will be fundamental to study mdx^5cv^-*JAG1*’s muscle satellite cells and their regenerative capacities. Also, since modulation of the Notch pathway has shown positive and deleterious effects in skeletal muscles (16, 19, 20), it will be essential to understand the precise biochemical mechanism downstream of *JAG1* overexpression (and supra-expression). Our proteomics dataset gave a glimpse into the beneficial mechanisms of increased expression of *JAG1* in dystrophic muscles. However, contractile proteins are the overwhelming majority of proteins in skeletal muscles, limiting the number of proteins found.

In conclusion, we show that muscle-specific *increased* JAG1 expression leads to increases in muscle size and force. This effect is noticeable over time, *so JAG1* overexpression continues to counterbalance the progressive consequences of dystrophin absence in the mouse. Finally, although our data support the exploration of the Notch pathway for DMD therapy, it must be done with care since harmful consequences of Notch modulation have been clearly described in humans (19, 20).

## Materials and Methods

### mdx^5cv^-Jag1 generation

All experimental procedures followed the Boston Children’s Hospital Animal Care and Use Committee (protocol #00001392). All strains used are on C57BL6/J background (JAX #000664). First, we crossed X-linked muscular dystrophy 5 (mdx^5cv^, JAX #002379) with R26-LSL-*JAG1* (JAX #030173) to establish mdx^5cv^ carrying human *JAG1* (mdx^5cv^; R26-LSL-*JAG1*). Next, we crossed MCK-Cre (JAX #006475) males with mdx^5cv^-Jag1 Cre^-^ females, establishing mdx^5cv^-*JAG1* Cre^+^. We studied male littermates Cre^+^ (mdx^5cv^-*JAG1*) and Cre^-^ (mdx^5cv^) carrying *JAG1 in* heterozygosis.

### Histology and immunofluorescence

TA muscles were dissected and snap-frozen in isopentane. Ten micrometers (μm) muscle cross-sections were obtained from frozen muscles using a Cryostat (Leica CM3050 S). Standard hematoxylin and eosin stain was performed. For immunostaining, frozen muscle cross-section slides were air-dried for 20 minutes. The slides were treated with blocking solution (1X PBS supplemented with 0.1% Tween-20 and 10% goat serum, #G9023 Sigma-Aldrich) for 1 hour at room temperature to avoid non-specific binding. Fiber typing was accomplished by histochemical staining for different myosin heavy chain isoforms (MyHC). Primary antibodies BA-D5-s (mouse IgG2b, dilution 1:50) for MyHC-slow, SC-71-s (mouse IgG1, dilution 1:50) for MyHC2A, and BF-F3-c (mouse IgM, dilution 1:50) for MyHC2B were obtained from the Developmental Studies Hybridoma Bank (DSHB). Unstained fibers were considered type 2X. Anti-laminin (anti-rabbit, #L9393, Sigma-Aldrich, dilution 1:500) was used to outline the fibers. Primary antibodies were diluted in blocking solution and incubated for two hours at 37°C. Sections were washed thrice for five minutes each with PBST (1X PBS with 0.1% Tween-20). Secondary antibodies DyLight 405 goat anti-mouse IgG2b (RRID: AB_2338801, Jackson ImmunoResearch), Alexa Fluor 594 goat anti-mouse IgG1 (RRID: AB_2338885, Jackson ImmunoResearch), Alexa Fluor 488 goat anti-mouse IgM (RRID: AB_2338849, Jackson ImmunoResearch), and Alexa Fluor 647 goat anti-rabbit IgG (H+L) (#A21244, Thermo Scientific) were diluted 1:200 in blocking solution and incubated for one hour at 37°C. Sections were washed thrice for five minutes with PBST, and the slides were mounted with Vectashield antifade mounting media (H-1000, Vector laboratories). An entire TA section was imaged using a 10x magnification objective on an Axio Imager.Z2 (Zeiss) microscope. Images were then analyzed using MyoSight macro for ImageJ (54). Non-stained fibers were considered type IIX fibers, and, as expected for TA muscles, type I fibers made up less than 1% of the total in both groups and were not considered.

For jagged-1 staining, sections on slides were fixed in ice-cold methanol for three minutes and washed in ice-cold PBS for one minute. Next, slides were incubated in blocking solution (PBS, 10% fetal bovine serum, and 0.1% Triton). Primary incubation was done overnight in a wet chamber at 4°C. Primary antibodies were AbCam 85763 (rabbit polyclonal) for jagged-1 and LS-C96142 (chicken) for laminin. On the following day, slides were washed in PBS for five minutes and incubated in secondary antibodies, anti-rabbit Alexa Fluor 488 and anti-chicken Alexa Fluor 647 (A21449). Finally, slides were washed in PBS and sealed with Vectashield with DAPI (Vector Laboratories: H-1200). Confocal images were taken using a Leica SP8 microscope.

For eMyHC, slides were fixed in 1:1 methanol and acetone for two minutes at -20°C, followed by thrice washes in PBS for three minutes. Next, slides were incubated in 5% goat serum in PBS for one hour. Primary incubation was done overnight in a wet chamber at 4°C with DSHB F1.652 (mouse monoclonal) for eMyHC and LS-C96142 (chicken) for laminin. On the following day, slides were washed four times for three minutes each in PBS and incubated with anti-mouse Alexa fluor 488 and anti-chicken Alexa Fluor 647 (A21449) for one hour. After an additional four washes of three minutes each, slides were sealed with Vectashield with DAPI (Vector Laboratories: H-1200). Images were made using a 20x magnification objective on an Axio Imager.Z2 (Zeiss) microscope.

### Western blot

Mechanical grinding of snap-frozen quadriceps muscles was performed with mortar and pestle in liquid nitrogen. Muscle powder was transferred to cold RIPA buffer (Boston BioProducts) containing protease inhibitors (cOmplete, mini, EDTA-free, Roche) and phosphatase inhibitors (PhosSTOP, Sigma-Aldrich). Tissue samples were then briefly sonicated, homogenized for 1 hour at 4°C, and centrifuged at 13,500 rpm for 30 mins at 4°C. The supernatant was collected, and protein concentration was determined using a Pierce BCA protein assay kit (Thermo Scientific). For SDS-PAGE, protein samples were denatured in Novex Tris-glycine SDS sample buffer (Invitrogen) containing 50 mM dithiothreitol (NuPAGE reducing agent, Invitrogen) and boiled for 8 mins at 90°C. Thirty micrograms of total protein were separated by electrophoresis on Novex 4-20% tris-glycine mini protein gels (Invitrogen) and transferred to 0.2-mm PVDF membrane (Invitrogen). The quality of protein transfer was checked with Ponceau stain (Thermo Scientific) before proceeding to Western blotting. Membranes were treated with 5% BSA in TBST buffer (tris-buffered saline containing 0.05% Tween-20) for one hour at room temperature to decrease non-specific binding. Membranes were then incubated with anti-jagged1 antibody (#ab300561, Abcam, dilution 1:1000) in blocking solution overnight at 4°C. Following three ten-minute-long washes in TBST buffer, membranes were probed with anti-rabbit IgG, HRP-linked secondary antibody (#7074S, Cell Signaling Technology) in blocking solution for two hours at room temperature. Membranes were then washed five times for ten minutes each in TBST buffer. Chemiluminescence detection was captured using Pierce ECL western blotting substrate (Thermo Scientific) and Biorad ChemiDoc system. The jagged-1 protein signal was analyzed using Image Lab software (Biorad) and normalized to the respective Ponceau signal.

### ex-vivo Physiology

The functional properties of the tibialis anterior (TA) were assessed using an in situ preparation. An anesthetized mouse (sodium pentobarbital, i.p. 80 mg/kg) was stabilized on a temperature-controlled platform (38°C, Aurora Scientific model 809), and the severed distal tendon of the TA was attached to the lever arm of a dual-model muscle lever system (Aurora Scientific model 305C-LR). Supramaximal 200 μs square-wave pulses were generated by a muscle stimulator (Aurora Scientific model 701) and delivered to platinum needle electrodes inserted behind the knee near the peroneal nerve. Muscle length was adjusted so that force to brief tetani was maximized. A minimum of 60 seconds separated all contractions to minimize fatigue. The force-frequency relationship was evaluated by stimulating the TA with trains of increasing frequency (from 10 up to 200 Hz). Five lengthening, or eccentric, contractions were used to assess the resistance of the TA to high-strain contractile activity. These contractions consisted of an initial 100 ms at a fixed length, allowing the preparation to attain peak tetanic force, and the remainder with the muscle lengthened by 20% of fiber length at 1.5 fiber lengths/s (assuming a fiber length to muscle length ratio of 0.60). The series of lengthening contractions were bracketed by fixed-end contractions for assessing pre- and post-lengthening tetanic force. The physiological cross-sectional area of the muscle was estimated as previously described (55).

### Data Analyses

Data analyses and visualization were performed using RStudio (R version 4.3.3) and the ggplot2 package. For comparisons consisting of one dependent variable (e.g., body mass, muscle mass, and peak force), normality was assessed using the Shapiro-Wilk test, and homogeneity of variances was evaluated with Levene’s test. Depending on the variance results, either a Student’s t-test (equal variances) or Welch’s t-test (unequal variances) was used for group comparisons. Results are presented as mean ± SEM, and the p-value is shown in each plot. A two-way ANOVA was used to test the effects of expression levels (baseline, increased, supra) and genotypes (mdx^5cv^ and mdx^5cv^-*JAG1*) on the variables when animals with supra-abundance of Jagged-1 were considered a separate group.

For non-linear regression modeling of force-frequency relationships and resistance to eccentric contraction-induced damage, a four-parameter log-logistic sigmoid function was used to fit data from different genotypes at three time points using the drm() function from the drc package for R. Genotype-specific models were compared to combined models using an F-test to evaluate whether separate genotype-specific curves provided a significantly better fit to the data than a single shared curve. Plots show mean ± SEM and fitted non-linear models (solid lines) with shaded 95% confidence intervals.

### Off-line basic pH reversed-phase (BPRP) fractionation

We fractionated the pooled TMT-labeled peptide (specifically, the unbound and wash from the phosphopeptide enrichment) via BPRP HPLC with an Agilent 1260 pump. Peptides were subjected to a 560 min linear gradient from 5% to 35% acetonitrile in 10 mM ammonium bicarbonate pH 8 at a flow rate of 0.25 mL/min over an Agilent 300Extend C18 column (3.5 μm particles, 2.1 mm ID and 25 cm long). The peptide mixture was fractionated into a total of 96 fractions, which were consolidated into 24 super-fractions and subjected to FAIMS-MS/MS. These fractions were subsequently acidified with 1% formic acid and vacuum centrifuged to near dryness. Each fraction was also desalted via StageTip, dried via vacuum centrifugation, and reconstituted in 5% acetonitrile, 5% formic acid for LC-MS/MS.

### Mass spectrometric data collection

Mass spectrometric data were collected on an Orbitrap Eclipse mass spectrometer coupled to a Vanquish Neo UHPLC. Approximately 1µg of peptide was separated at a flow rate of 450 nL/min on a 100 µm capillary column that was packed with 35 cm of Accucore 150 resin (2.6 μm, 150Å; ThermoFisher Scientific). The scan sequence began with an MS1 spectrum (Orbitrap analysis, resolution 60,000, 350-1350 Th, automatic gain control (AGC) target is set to 100%, maximum injection time set to 50ms). Data were acquired 90 minutes per fraction. The hrMS2 stage consisted of fragmentation by higher energy collisional dissociation (HCD, normalized collision energy 36%) and analysis using the Orbitrap (AGC 200%, maximum injection time 120ms, isolation window 0.6 Th, resolution 30,000 Turbo TMT). Data were acquired using the FAIMSpro interface the dispersion voltage (DV) set to 5,000V, the compensation voltages (CVs) were set at -30V, -50V, and -60V or -40V, -60V, and -70V. The TopSpeed parameter was set at 1 sec per CV.

### Mass spectrometric data analysis

Mass spectra were processed using a Comet-based in-house software pipeline. MS spectra were converted to mzXML using a modified version of ReAdW.exe. Database searching included all entries from the human UniProt database, which was concatenated with a reverse database composed of all protein sequences in reversed order (56, 57). Searches were performed using a 50 ppm precursor ion tolerance. Product ion tolerance was set to 0.03 Th. Carbamidomethylation of cysteine residues (+57.0215Da) were set as static modifications, while oxidation of methionine residues (+15.9949 Da) was set as a variable modification.

Peptide spectral matches (PSMs) were altered to a 1% FDR. PSM filtering was performed using a linear discriminant analysis, as described previously (58), while considering the following parameters: XCorr, ΔCn, missed cleavages, peptide length, charge state, and precursor mass accuracy. Peptide-spectral matches were identified, quantified, and collapsed to a 1% FDR and then further collapsed to a final protein-level FDR of 1%. Furthermore, protein assembly was guided by principles of parsimony to produce the smallest set of proteins necessary to account for all observed peptides. Spectral counts were then extracted and the data were subsequently analyzed.

### Proteomics Data analysis

Spectra were converted to mzXML via MSconvert (59). Database searching included all entries from human UniProt reference Database (downloaded: January 2024). The database was concatenated with one composed of all protein sequences for that database in the reversed order. Searches were performed using a 50-ppm precursor ion tolerance for total protein level profiling. The product ion tolerance was set to 0.03 Da. These wide mass tolerance windows were chosen to maximize sensitivity in conjunction with Comet searches and linear discriminant analysis (58, 60). TMTpro labels on lysine residues and peptide N-termini +304.207 Da), as well as carbamidomethylation of cysteine residues (+57.021 Da) were set as static modifications, while oxidation of methionine residues (+15.995 Da) was set as a variable modification. Peptide-spectrum matches (PSMs) were adjusted to a 1% false discovery rate (FDR) (56, 57). PSM filtering was performed using a linear discriminant analysis, as described previously (58) and then assembled further to a final protein-level FDR of 1% (56). Proteins were quantified by summing reporter ion counts across all matching PSMs, also as described previously (61). Reporter ion intensities were adjusted to correct for the isotopic impurities of the different TMTpro reagents according to manufacturer specifications. The signal-to-noise (S/N) measurements of peptides assigned to each protein were summed and these values were normalized so that the sum of the signal for all proteins in each channel was equivalent to account for equal protein loading. Finally, each protein abundance measurement was scaled, such that the summed signal-to-noise for that protein across all channels equals 100, thereby generating a relative abundance (RA) measurement.

## Acknowledgments

This publication is based on research supported by the MDA Award to FSL (https://doi.org/10.55762/MDA.1291397.pc.gr.198128). LMK and FSL received support from Defeat Duchenne Canada. MRL was supported by an AFM-Telethon postdoctoral fellowship (#21904) for this work. MRL is also a recipient of a Muscular Dystrophy Association grant (MDA 962927, DOI: 10.55762/pc.gr.157029), a NIH career development grant (K99AR080197), and a junior faculty career development grant from the Office of Faculty Development at Boston Children’s Hospital. TYZ was funded by American Heart Association postdoctoral fellowship (AHA18POST34070039) and NCI grant (R00CA273170). This work was supported by National Institutes of Health (R01AR064300 and gm067945). Funding from the Bernard F. and Alva B. Gimbel Foundation to LMK supported FSL. LMK received support from a IDDRC grant (P50HD105351). FSL was supported by CNPq (404161/2019-7).

## References

1. N. M. Vieira, et al., Jagged 1 Rescues the Duchenne Muscular Dystrophy Phenotype. Cell 163, 1204–1213 (2015).

2. M. Koenig, et al., Complete cloning of the Duchenne muscular dystrophy (DMD) cDNA and preliminary genomic organization of the DMD gene in normal and affected individuals. Cell 50, 509–517 (1987).

3. E. P. Hoffman, R. H. Brown Jr., L. M. Kunkel, Dystrophin: the protein product of the Duchenne muscular dystrophy locus. Cell 51, 919–928 (1987).

4. A. E. Emery, Population frequencies of inherited neuromuscular diseases--a world survey. Neuromuscul Disord 1, 19–29 (1991).

5. J. Broomfield, M. Hill, M. Guglieri, M. Crowther, K. Abrams, Life Expectancy in Duchenne Muscular Dystrophy. Neurology 97, e2304 LP–e2314 (2021).

6. E. Pegoraro, et al., SPP1 genotype is a determinant of disease severity in Duchenne muscular dystrophy. Neurology 76, 219–226 (2011).

7. A. Heydemann, et al., Latent TGF-beta-binding protein 4 modifies muscular dystrophy in mice. J Clin Invest 119, 3703–3712 (2009).

8. A. R. Demonbreun, et al., Anti-latent TGFβ binding protein 4 antibody improves muscle function and reduces muscle fibrosis in muscular dystrophy. Sci Transl Med 13, eabf0376 (2021).

9. K. M. Flanigan, et al., LTBP4 genotype predicts age of ambulatory loss in Duchenne muscular dystrophy. Ann Neurol 73, 481–488 (2013).

10. M. W. Hogarth, et al., Evidence for ACTN3 as a genetic modifier of Duchenne muscular dystrophy. Nat Commun 8, 14143 (2017).

11. K. M. Flanigan, et al., A genome-wide association analysis of loss of ambulation in dystrophinopathy patients suggests multiple candidate modifiers of disease severity. Eur J Hum Genet 31, 663–673 (2023).

12. S. Bogdanovich, et al., Functional improvement of dystrophic muscle by myostatin blockade. Nature 420, 418–421 (2002).

13. R. M. Hightower, M. S. Alexander, Genetic modifiers of Duchenne and facioscapulohumeral muscular dystrophies. Muscle Nerve 57, 6–15 (2018).

14. M. Zatz, et al., A normal life without muscle dystrophin. Neuromuscul Disord 25, 371–374 (2015).

15. E. Zucconi, et al., Ringo: discordance between the molecular and clinical manifestation in a golden retriever muscular dystrophy dog. Neuromuscul Disord 20, 64–70 (2010).

16. T. Kitamoto, K. Hanaoka, Notch3 null mutation in mice causes muscle hyperplasia by repetitive muscle regeneration. Stem Cells 28, 2205–2216 (2010).

17. P. Bi, et al., Stage-specific effects of Notch activation during skeletal myogenesis. Elife 5 (2016).

18. X. Wei, et al., Neurofibromin 1 controls metabolic balance and Notch-dependent quiescence of murine juvenile myogenic progenitors. Nat Commun 15, 1393 (2024).

19. E. Servián-Morilla, et al., A POGLUT1 mutation causes a muscular dystrophy with reduced Notch signaling and satellite cell loss. EMBO Mol Med 8, 1289–1309 (2016).

20. S. Coppens, et al., A form of muscular dystrophy associated with pathogenic variants in JAG2. Am J Hum Genet 108, 840–856 (2021).

21. I. Danko, V. Chapman, J. A. Wolff, The frequency of revertants in mdx mouse genetic models for duchenne muscular dystrophy. Pediatr Res 32, 128–131 (1992).

22. Q. Su, et al., Jagged1 upregulation in prostate epithelial cells promotes formation of reactive stroma in the Pten null mouse model for prostate cancer. Oncogene 36, 618– 627 (2017).

23. J. C. Brüning, et al., A muscle-specific insulin receptor knockout exhibits features of the metabolic syndrome of NIDDM without altering glucose tolerance. Mol Cell 2, 559–569 (1998).

24. D. Bansal, et al., Defective membrane repair in dysferlin-deficient muscular dystrophy. Nature 423, 168–172 (2003).

25. A. K. Lucio-Eterovic, et al., Role for the nuclear receptor-binding SET domain protein 1 (NSD1) methyltransferase in coordinating lysine 36 methylation at histone 3 with RNA polymerase II function. Proceedings of the National Academy of Sciences 107, 16952– 16957 (2010).

26. N. Huang, et al., Two distinct nuclear receptor interaction domains in NSD1, a novel SET protein that exhibits characteristics of both corepressors and coactivators. EMBO J 17, 3398–3412 (1998).

27. T. Baubec, et al., Genomic profiling of DNA methyltransferases reveals a role for DNMT3B in genic methylation. Nature 520, 243–247 (2015).

28. H. Watanabe, et al., DNA methylation analysis of multiple imprinted DMRs in Sotos syndrome reveals IGF2-DMR0 as a DNA methylation-dependent, P0 promoter-specific enhancer. The FASEB Journal 34, 960–973 (2020).

29. M. L. van den Boogaard, et al., Mutations in DNMT3B Modify Epigenetic Repression of the D4Z4 Repeat and the Penetrance of Facioscapulohumeral Dystrophy. Am J Hum Genet 98, 1020–1029 (2016).

30. L. F. Bouwman, et al., Dnmt3b regulates DUX4 expression in a tissue-dependent manner in transgenic D4Z4 mice. Skelet Muscle 10, 27 (2020).

31. T. Saito, et al., Coagulation and fibrinolysis disorder in muscular dystrophy. Muscle Nerve 24, 399–402 (2001).

32. P. H. Berman, M. A. Nigro, M. B. Harris, F. A. Oski, Activation of fibrinolysis in muscular dystrophy. Arch Neurol 29, 65–66 (1973).

33. A. M. S. Yamashita, et al., Effect of Notch1 signaling on muscle engraftment and maturation from pluripotent stem cells. Stem Cell Reports 20 (2025).

34. C. Webster, L. Silberstein, A. P. Hays, H. M. Blau, Fast muscle fibers are preferentially affected in Duchenne muscular dystrophy. Cell 52, 503–513 (1988).

35. J. Talbot, L. Maves, Skeletal muscle fiber type: using insights from muscle developmental biology to dissect targets for susceptibility and resistance to muscle disease. Wiley Interdiscip Rev Dev Biol 5, 518–534 (2016).

36. M. Pedemonte, C. Sandri, S. Schiaffino, C. Minetti, Early decrease of IIx myosin heavy chain transcripts in Duchenne muscular dystrophy. Biochem Biophys Res Commun 255, 466–469 (1999).

37. J. F. Marini, et al., Expression of myosin heavy chain isoforms in Duchenne muscular dystrophy patients and carriers. Neuromuscul Disord 1, 397–409 (1991).

38. L. T. Krebs, et al., Characterization of Notch3-deficient mice: normal embryonic development and absence of genetic interactions with a Notch1 mutation. Genesis 37, 139–143 (2003).

39. N. M. Vieira, et al., Repression of phosphatidylinositol transfer protein alpha ameliorates the pathology of Duchenne muscular dystrophy. Proc Natl Acad Sci U S A 114, 6080–6085 (2017).

40. M. R. Lambert, et al., PDE10A Inhibition Reduces the Manifestation of Pathology in DMD Zebrafish and Represses the Genetic Modifier PITPNA. Mol Ther 29, 1086–1101 (2021).

41. A. C. McPherron, A. M. Lawler, S. J. Lee, Regulation of skeletal muscle mass in mice by a new TGF-beta superfamily member. Nature 387, 83–90 (1997).

42. S. Girgenrath, K. Song, L.-A. Whittemore, Loss of myostatin expression alters fiber-type distribution and expression of myosin heavy chain isoforms in slow- and fast-type skeletal muscle. Muscle Nerve 31, 34–40 (2005).

43. R. Bashir, et al., A gene related to Caenorhabditis elegans spermatogenesis factor fer-1 is mutated in limb-girdle muscular dystrophy type 2B. Nat Genet 20, 37–42 (1998).

44. J. Liu, et al., Dysferlin, a novel skeletal muscle gene, is mutated in Miyoshi myopathy and limb girdle muscular dystrophy. Nat Genet 20, 31–36 (1998).

45. R. Han, E. P. Rader, J. R. Levy, D. Bansal, K. P. Campbell, Dystrophin deficiency exacerbates skeletal muscle pathology in dysferlin-null mice. Skelet Muscle 1, 35 (2011).

46. L. E. Glover, et al., Dysferlin overexpression in skeletal muscle produces a progressive myopathy. Ann Neurol 67, 384–393 (2010).

47. G.-Y. Jeong, et al., NSD3-Induced Methylation of H3K36 Activates NOTCH Signaling to Drive Breast Tumor Initiation and Metastatic Progression. Cancer Res 81, 77–90 (2021).

48. Y. Wen, A. P. Alimov, J. J. McCarthy, Ribosome Biogenesis is Necessary for Skeletal Muscle Hypertrophy. Exerc Sport Sci Rev 44, 110–115 (2016).

49. A. Lu, et al., Rapid depletion of muscle progenitor cells in dystrophic mdx/utrophin−/− mice. Hum Mol Genet 23, 4786–4800 (2014).

50. Z. Yablonka-Reuveni, R. Seger, A. J. Rivera, Fibroblast growth factor promotes recruitment of skeletal muscle satellite cells in young and old rats. J Histochem Cytochem 47, 23–42 (1999).

51. Z. Yablonka-Reuveni, A. J. Rivera, Proliferative Dynamics and the Role of FGF2 During Myogenesis of Rat Satellite Cells on Isolated Fibers. Basic Appl Myol 7, 189–202 (1997).

52. N. Li, et al., Identification of hub genes and therapeutic siRNAs to develop novel adjunctive therapy for Duchenne muscular dystrophy. BMC Musculoskelet Disord 25, 386 (2024).

53. I. Spector, et al., The Involvement of Collagen Triple Helix Repeat Containing 1 in Muscular Dystrophies. Am J Pathol 182, 905–916 (2013).

54. L. W. Babcock, A. D. Hanna, N. H. Agha, S. L. Hamilton, MyoSight-semi-automated image analysis of skeletal muscle cross sections. Skelet Muscle 10, 33 (2020).

55. M. S. Alexander, et al., MicroRNA-486-dependent modulation of DOCK3/PTEN/AKT signaling pathways improves muscular dystrophy-associated symptoms. J Clin Invest 124, 2651–2667 (2014).

56. J. E. Elias, S. P. Gygi, Target-decoy search strategy for increased confidence in large-scale protein identifications by mass spectrometry. Nat Methods 4, 207–214 (2007).

57. J. E. Elias, S. P. Gygi, Target-decoy search strategy for mass spectrometry-based proteomics. Methods Mol Biol 604, 55–71 (2010).

58. E. L. Huttlin, et al., A tissue-specific atlas of mouse protein phosphorylation and expression. Cell 143, 1174–1189 (2010).

59. M. C. Chambers, et al., A cross-platform toolkit for mass spectrometry and proteomics. Nat Biotechnol [Preprint] (2012).

60. S. A. Beausoleil, J. Villén, S. A. Gerber, J. Rush, S. P. Gygi, A probability-based approach for high-throughput protein phosphorylation analysis and site localization. Nat Biotechnol 24, 1285–1292 (2006).

61. G. C. McAlister, et al., Increasing the multiplexing capacity of TMTs using reporter ion isotopologues with isobaric masses. Anal Chem 84, 7469–7478 (2012).

