## Supplemental Material for "Muscle-specific increased expression of *JAG1* improves skeletal muscle phenotype in dystrophin-deficient mice"

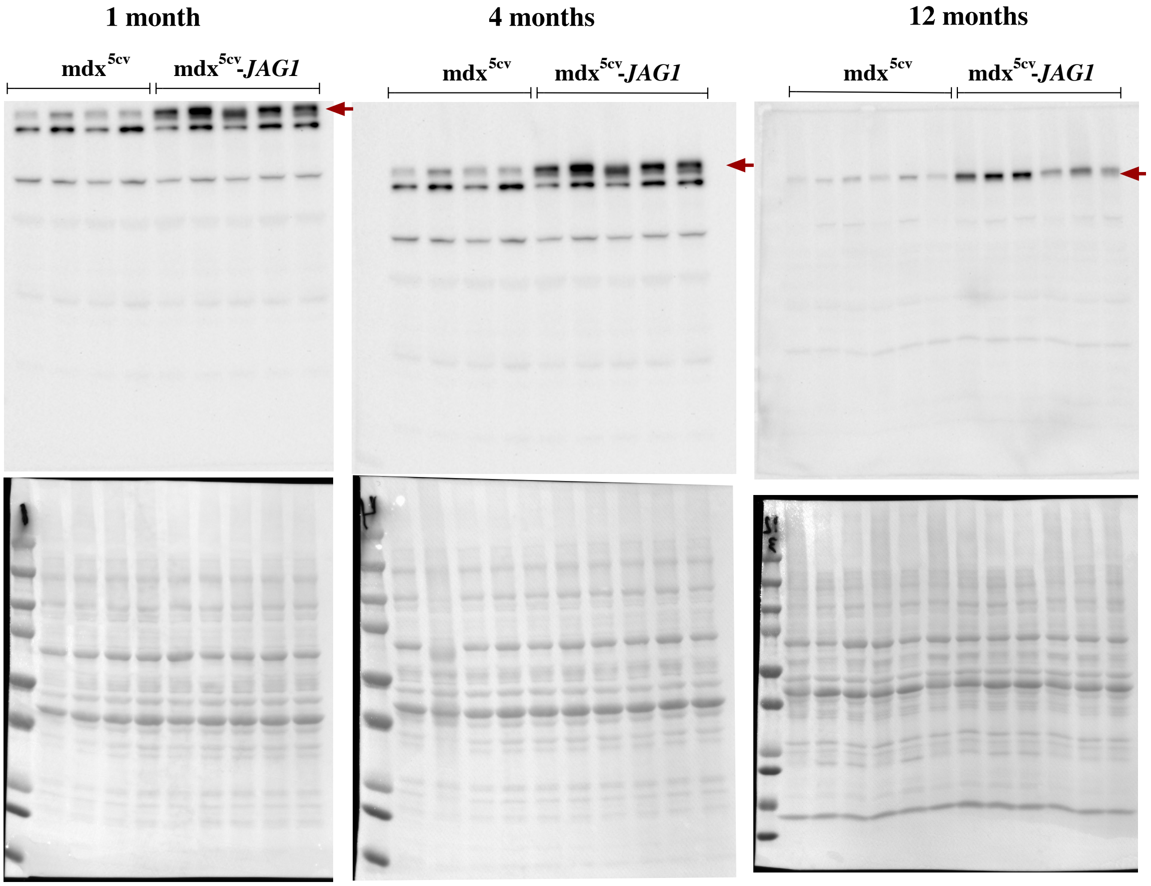


**Supplemental Figure 1 - Membranes used for western blots in figure 1.** From left to right, images of the entire membrane at one, four, and twelve months. Top row shows membranes stained for Jagged-1 and bottom row shows membranes stained with Ponceau for load control. Red arrows point to Jagged-1 bands.

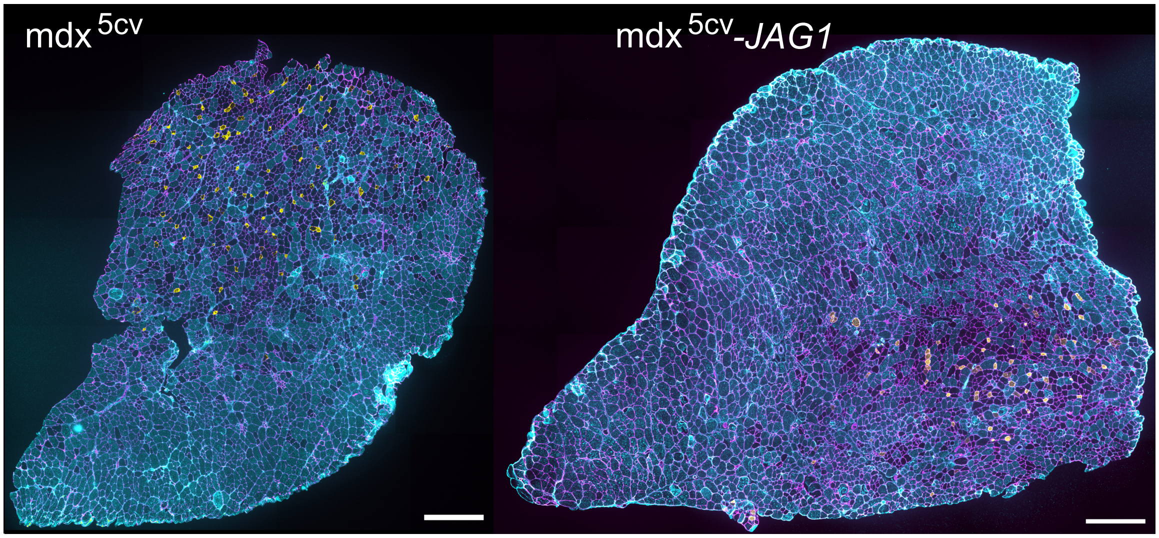


**Supplemental Figure 2 - Comparison of entire cross-sections of one year-old mdx$^{5cv}$ and mdx$^{5cv}$-JAG1.** Overlay of stained sections obtained from the thickest portion of the TA muscle. Sections were stained for Laminin (Magenta), Myosin heavy chain type I (not shown), type IIA (yellow), and IIB (Cyan). Scale bars are 100 µm. This figure is related to Figure 3 in the main text.

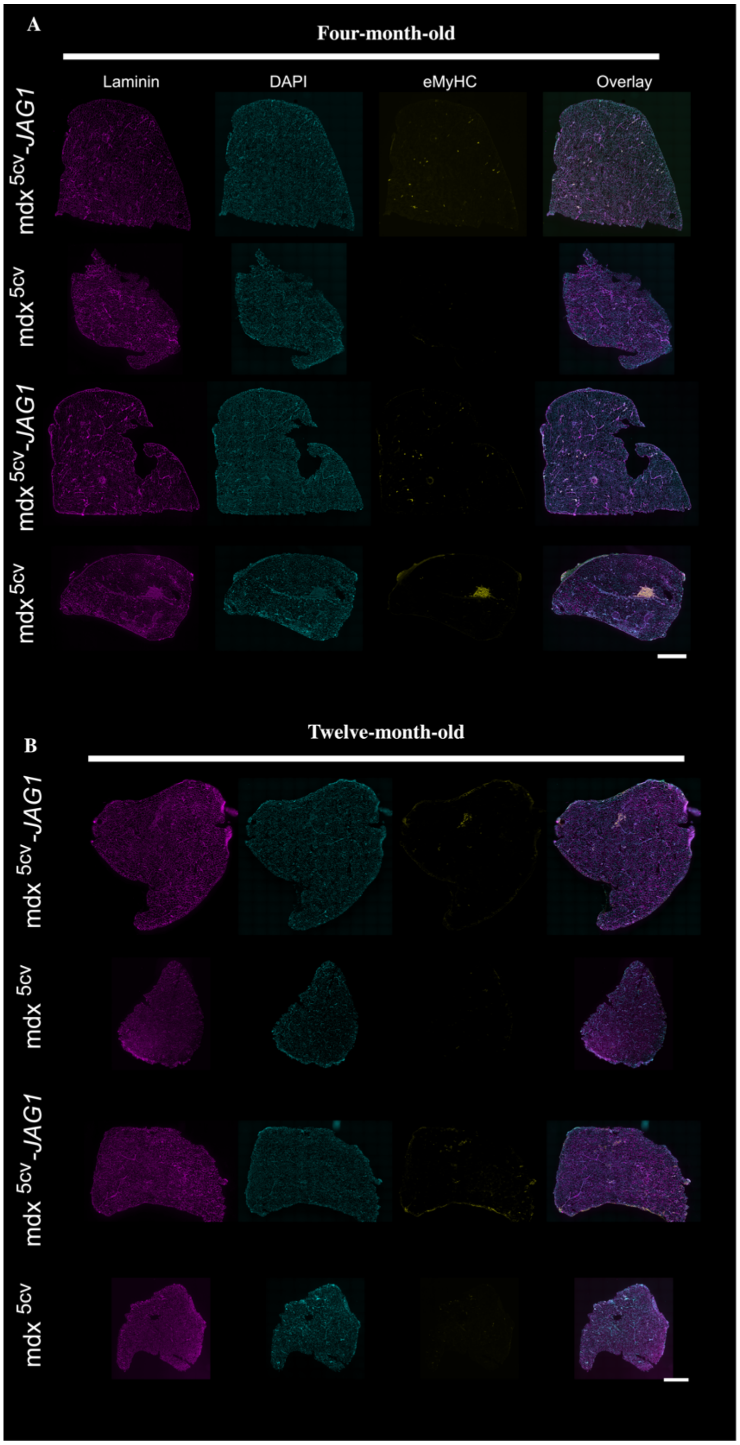


**Supplemental Figure 3 - Embryonic Myosin Heavy Chain staining of (A) four- and (B) twelve-month-old sections.** TA sections. From left to right, the figure shows laminin staining in magenta, nuclear DNA staining with DAPI in cyan, embryonic myosin heavy chain (eMyHC) in yellow, and the overlay of all channels. From top to bottom, the figure intercalates $mdx^{5cv}$-JAG1 and $mdx^{5cv}$ two times. The scale bar is 500 μm. This figure is related to Figure 4 in the main text.

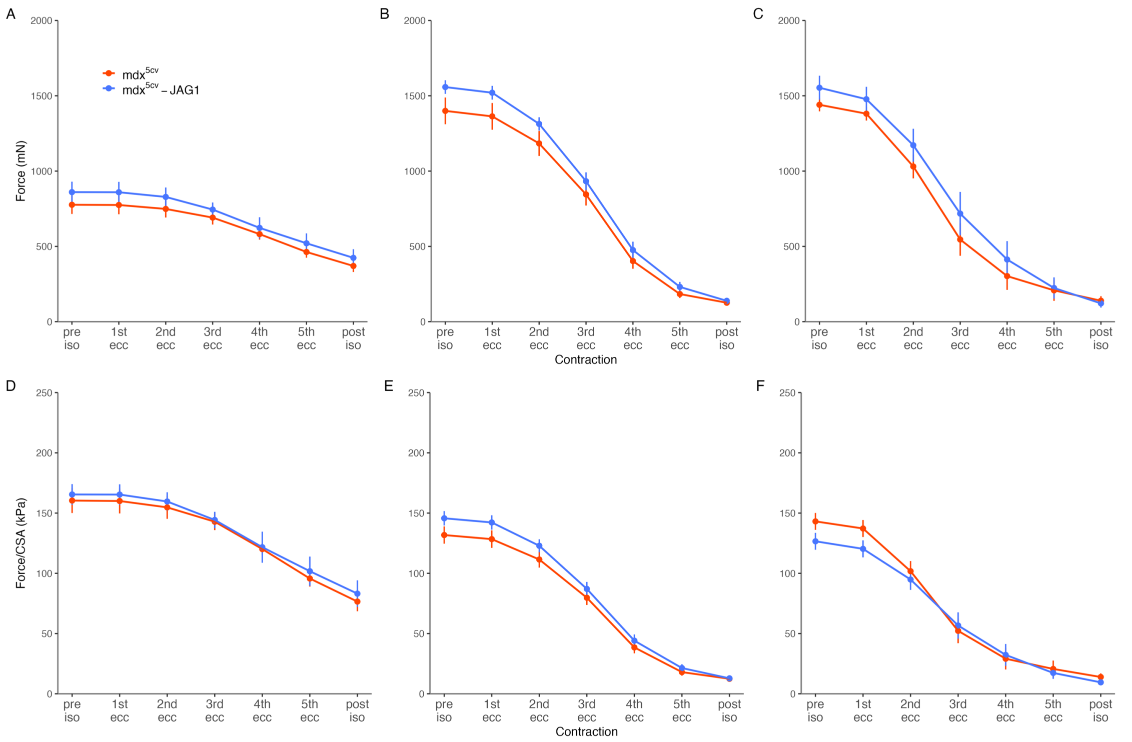


**Supplemental Figure 4 – Pre- and post-isometirc force and the isometric force measured during the preliminary isometric phase of each eccentric contraction. (A-C)** Absolute force measured during the eccentric contraction protocol. **(D-F)** Force/CSA measured during the eccentric contraction protocol. Plots show mean ±SEM for each contraction.

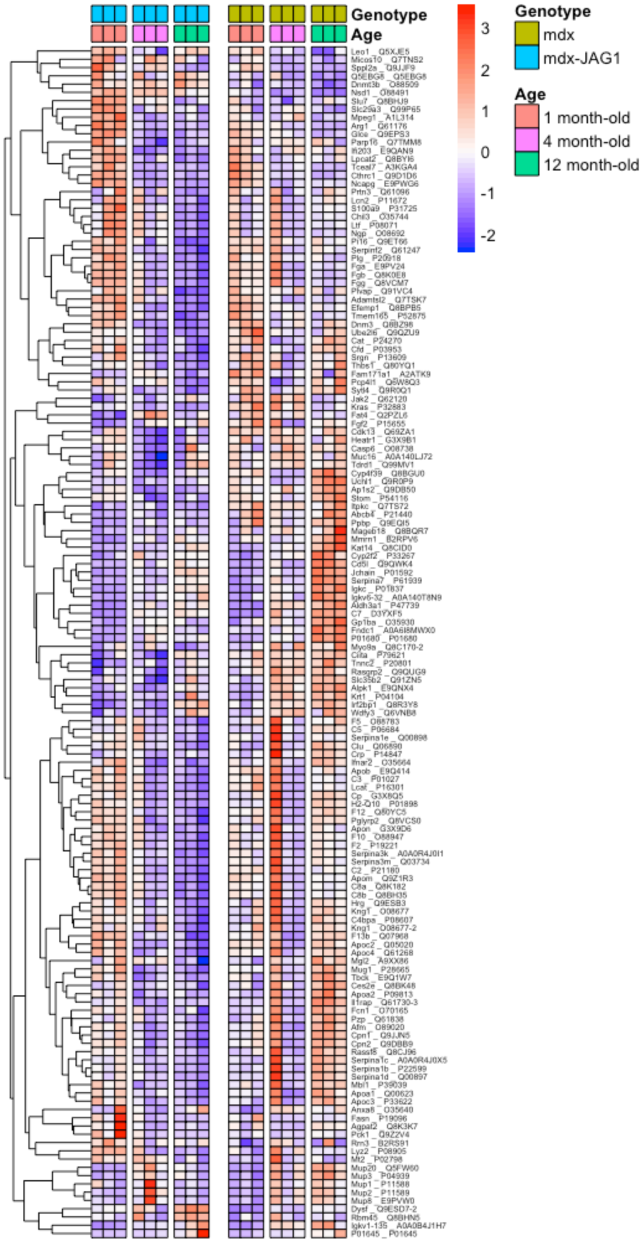


**Supplemental Figure 5 – Time course response of all proteins found with differential abundance at a given time point.** Heat map showing the Z-score normalized per line.

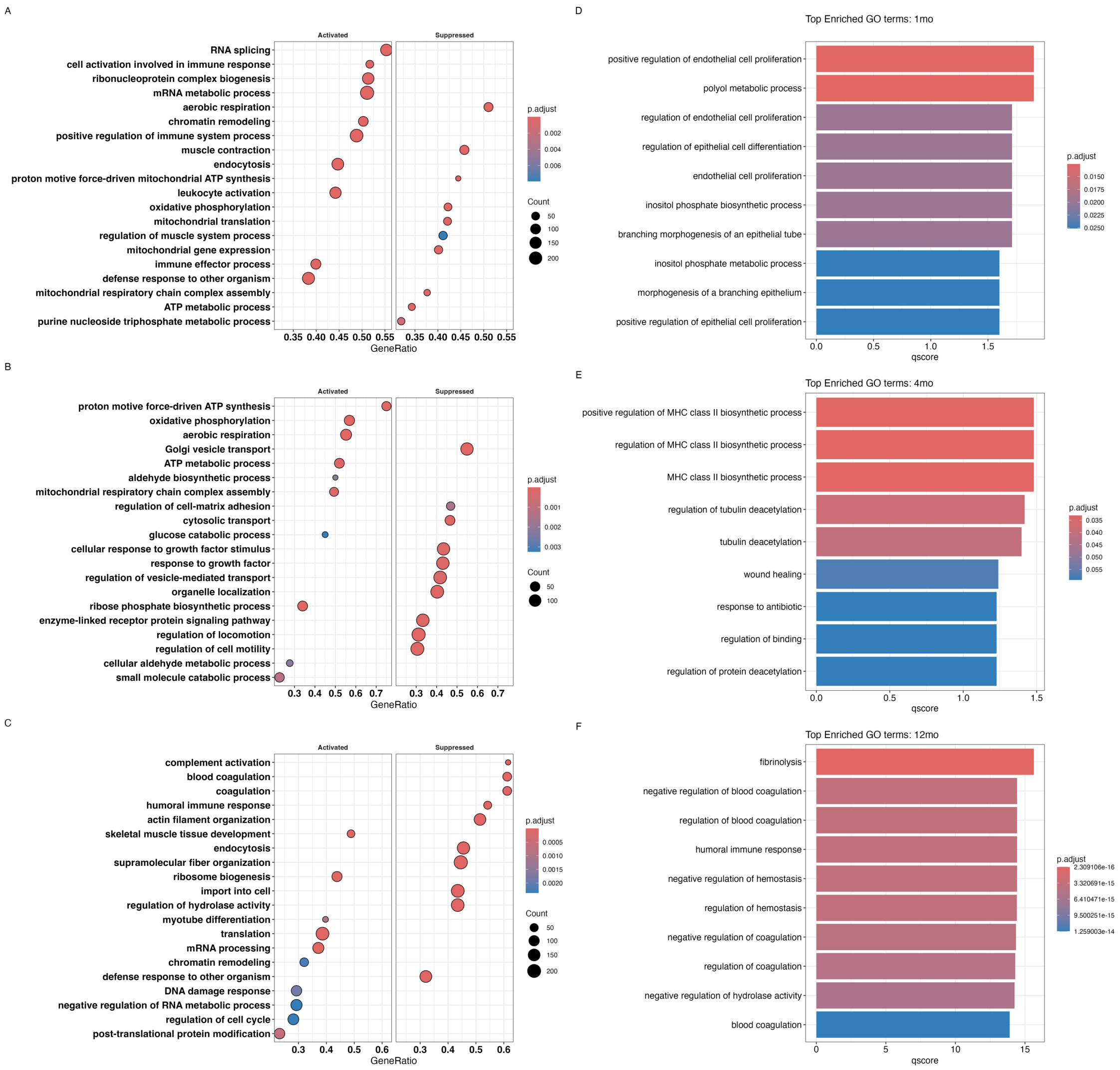


**Supplemental Figure 6 – (A-C)** Top enriched Gene Ontology terms from Gene Set Enrichment Analysis. From top to bottom, one-, four-, and twelve-month-old analyses. Left panel shows active pathways and right panel shows suppressed ones. **(D-F)** Top enriched Gene Ontology terms from over-representation analysis (ORA).

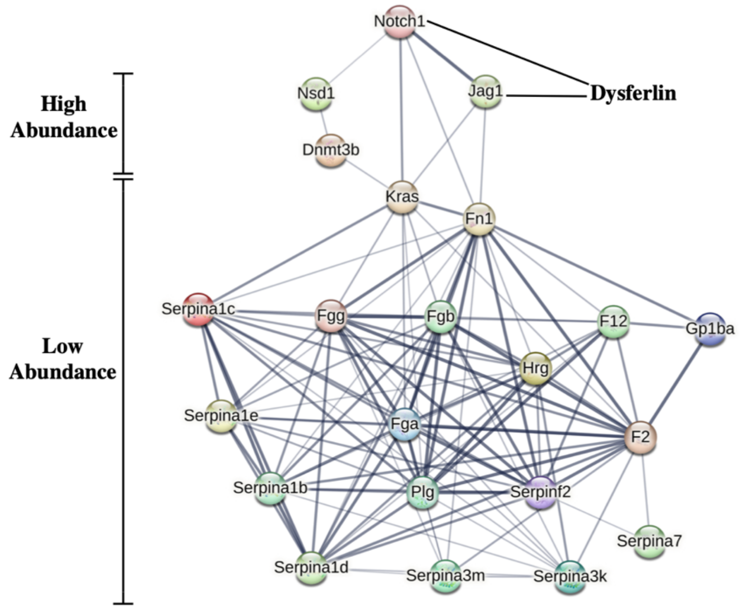


**Supplemental Figure 7 -** STRING map showing associations among proteins showing different abundance at the one-year time point. Dysferlin was manually added based on our findings and (17). Notch1 one was added as an example of Notch receptor that interacts with Jagged-1.
